# Seed Microbiota Diversity and Culture Collection of Four Major Crops Covering Different Genotypes and Production Modes

**DOI:** 10.64898/2026.04.29.721552

**Authors:** Marie Simonin, Natalia Guschinskaya, Muriel Marchi, Coralie Marais, Anne Préveaux, Martial Briand, Nathan Kavunu, Anaïs Bosc-Bierne, Laurine Labourgade, Cécile Dutrieux, Agathe Brault, Sophie Rolland, Claude-Emmanuel Koutouan, Perrine Portier, Mathilde Causse, Thierry Langin, Nathalie Nesi, Nicolas WG Chen, Alain Sarniguet, Matthieu Barret

## Abstract

Seed microbiota play a crucial role in plant health and development, yet remain understudied compared to other plant-associated microbial communities. This study aimed to characterize seed microbiota diversity across four major crops (common bean, rapeseed, tomato, and wheat) and establish a comprehensive strain collection of seed-borne microorganisms (bacteria and fungi). We employed a combination of culture-dependent and culture-independent approaches to analyze 68 seed samples representing diverse genotypes and production modes.

Our results revealed highly variable seed microbiota, with bacterial colonization ranging from 10 to 100 million bacterial CFUs per gram of seeds, and microbial richness varying from 4 to 351 bacterial and 16 to 138 fungal amplicon sequence variants (ASVs) per sample. Both plant genotype and production mode significantly influenced microbiota composition, with each seed sample produced harboring a distinct microbial assemblage. Interestingly, seeds produced in confined environments exhibited lower bacterial colonization but higher microbial richness compared to field-produced seeds. We observed divergent ecological drivers shaping bacterial and fungal communities. Bacterial assemblages were more host-specific and variable, while fungal communities showed greater stability and a substantial core microbiome shared across plant species. Our culturomics approach yielded a collection of 2,510 bacterial and 837 fungal isolates, representing 10-21% of the seed microbiota diversity detected by metabarcoding and the majority of the prevalent and abundant taxa. Notably, 44-60% of cultured bacterial isolates were not detected by metabarcoding, highlighting the complementary nature of these approaches to detect rare or under amplified taxa in PCR. This study provides insights into the complexity and variability of seed microbiota across different crops and production conditions. Our findings emphasize the importance of combining culturomics and sequencing methods for comprehensive characterization of seed microbiota to uncover the potential of seed-borne microorganisms as bioinoculants for sustainable agriculture.

## INTRODUCTION

Seeds represent a cornerstone of agriculture and for the maintenance of plant biodiversity (Leck *et al*. 2008, Reeves, Thomas, and Ramsay 2016). Most seeds host a microbiota composed of bacterial, archaeal and fungal taxa which can colonize various habitats from the surface of the embryo to the seed coat (Shade, Jacques, and Barret 2017, Nelson 2018). Comparatively to other plant compartments, seed microbiota have received less attention despite its potential importance for crop establishment and disease resistance (Bziuk *et al*. 2021, Matsumoto *et al*. 2021, Rétif *et al*. 2023). The published studies generally cover only one or few plant genotypes and production sites and we currently lack data to extensively characterize seed microbiota variability between diverse productive environments.

Seed microbiota is the starting point of microbiota assembly providing a primary inoculum to the germinating seed and growing plant (Shade, Jacques, and Barret 2017, Johnston-Monje, Gutiérrez, and Lopez-Lavalle 2021). Seed microbial communities are less diverse and abundant than other plant compartments with a specific assemblage of microorganisms (Leff *et al*. 2017, Nelson 2018). Seed microbiota represents a reservoir of plant-adapted and potentially vertically transmitted taxa with beneficial effects (Bergna *et al*. 2018, Kim *et al*. 2022, Zhang X *et al*. 2022, Hu *et al*. 2024).

Research on alternative solutions to pesticides and synthetic fertilizers is rising to mitigate their detrimental environmental impacts and following societal demands. One avenue being the use of microbial inoculants and microbiome engineering which can have multiple plant beneficial effects for stress tolerance and growth promotion (Elnahal *et al*. 2022). To implement these biostimulant and biocontrol methods to the field, seeds represent an ideal vector based on biopriming or seed coating methods (O’Callaghan, Ballard, and Wright 2022, Joubert *et al*. 2025). To develop efficient inoculation strategies with adapted microbes to this habitat, seed microbiota represent an untapped reservoir of microorganisms to explore. Still to date, few extensive culturomics approaches have been attempted to obtain large microbial strain collections from seeds (Matsumoto *et al*. 2021, Marchi *et al*. 2026). Hence the beneficial functions and transmission potential to growing plants of seed-borne microorganisms is underused, and current commercial bioinoculants are often isolated from roots with mixed results on the field (Kaminsky *et al*. 2019, Russ *et al*. 2023).

In this study, we aimed to extend knowledge on seed microbiota diversity of four major crops (common bean, rapeseed, tomato, wheat) and build a large strain collection of bacteria and fungi isolated from seeds. These plant species cover a diversity of types of fruits (pod, silique, berry, caryopsis), seeds (exalbuminous, albuminous) and pollination (auto or allogamous). Our specific objectives were to: 1) evaluate the effect of plant genotypes and production modes on seed microbiota diversity, taxonomy and community sizes and 2) isolate a large diversity of bacterial and fungal strains and assess the representativeness of the strain collection compared to the initial microbial diversity of the seed samples. This strain collection will represent a valuable resource for future fundamental synthetic community studies and the identification of bioinoculants in agriculture.

## MATERIALS AND METHODS

### Production of seed material

Seeds of the four crops were produced in 2020 (bean, tomato, rapeseed) and 2021 (wheat) at different experimental stations in France (Table 1). For each crop species, seeds of 8 to 10 genotypes were selected to be representative of the genetic diversity cultivated in France. These genotypes were produced in two different production environments (sites or culture conditions), depending on the crop considered: bean seeds were produced in the field at two distant locations in France (Anjou and Gascogne regions), tomato seeds were produced in the field or in a greenhouse in Provence region, wheat seeds in greenhouse or climatic chamber conditions in Auvergne region and rapeseed seeds were obtained in either open or self-pollinated conditions in Brittany region. The self-pollinated condition consisted in a treatment using fly pollination in insect proof cages to ensure autogamy and therefore produce genetically homogeneous lines. A total of 68 seed samples were produced for the study: 16 bean samples (8 genotypes x 2 production sites), 20 rapeseed samples (10 genotypes x 2 production modes), 16 tomato samples (8 genotypes x 2 production modes), 16 wheat samples (8 genotypes x 2 production modes).

**Table 1:** Information regarding the seed material produced for this study.

| Crop | Production sites | Culture condition | Genotypes |
| --- | --- | --- | --- |
| Common Bean<br>(*Phaseolus vulgaris* L.) | FNAMS, Loire Authion, Anjou<br>(47.470377, -0.394989) | Field | Caprice, Contender, Deezer, Facila, Flavert, Linex, Vanilla, Vezer |
|  | FNAMS, Condom, Gascogne<br>(43.956065, 0.3921756470) | Field |  |
| Rapeseed<br>(*Brassica napus* L.) | IGEPP, Le Rheu, Bretagne<br>(48.13916, -1.799776) | Open Field | Astrid, Aviso, Boston, Gross Luesewitzer, Jupiter, Lipton, Major, Milena, Mohican, Ramses |
|  | IGEPP, Le Rheu, Bretagne | Self-pollinated Field |  |
| Tomato<br>(*Solanum lycopersicum* L.) | GAFL, Avignon, Provence | Field | Cervil, Criollo, Ferum, LA0147, LA1420, Levovil, Plov24A, Stupicke PolniRane |
|  | GAFL, Avignon, Provence | Greenhouse |  |
| Winter Wheat<br>(*Triticum aestivum*) | GDEC, Clermont Ferrand, Auvergne | Greenhouse | Apache, Barok, Illico, Julius, Recital, Renan, Rubisko, Sy Mathis |
|  | GDEC, Clermont Ferrand, Auvergne | Climatic Chamber |  |

Our main objective was to produce diverse seed materials for microbiota analysis and culturomics for each crop, and not to compare crop species between them. Thus, data and statistical analyses were generally conducted for each crop species separately as the production sites and culture conditions were not comparable.

### Sample preparation for seed microbiota characterization and isolation

Seeds were soaked in a solution of Phosphate Buffered Saline including 0.05% of Tween 20 (Fisher Scientific, Waltham, MA, USA) to collect both seed epiphytic and endophytic microorganisms following International Seed Testing Association (ISTA) guidelines. Three biological repetitions per seed sample (i.e. a genotype for a given production mode) were used for tomato (1g, approx. 360 seeds), wheat (4g, approx. 100 seeds), rapeseed (2g, approx. 400 seeds) and bean (3 to 19 g, approx. 30 seeds).

Maceration volumes of seed samples were different for each crop because of their different sizes and imbibition capacity: 4, 8, and 4 mL for tomato, wheat, and rapeseed respectively. Given the large size variation of bean seeds, the maceration volume was 2 mL/g of seeds (ranging from 6 to 38 mL). Seed maceration was conducted under agitation at 6°C for 2h30 for rapeseed, and 16h overnight for the 3 other crop species. After maceration, the tomato and wheat seeds were placed in a stomacher machine for 3 minutes. These soaking suspensions were then either used directly for microbial isolation (see below) or DNA extraction and metabarcoding analysis. Bacterial cultivable community size was assessed for each seed sample by serial-dilution and plating soaking suspensions on 1/10 strength tryptic soy agar medium, (TSA10, Oxoid, Basingstoke, UK) supplemented with cycloheximide (50 μg/mL, Sigma-Aldrich, Saint-Louis, MO, USA). Bacterial colony forming units (CFU) were counted after 7 days at 20°C. For DNA extraction, 1 mL of soaking suspensions was centrifuged (4,000 × g, 10 min, 4°C), supernatants were discarded, and the pellet was stored at -80°C.

In addition to seeds, for microbial isolation only, we also analyzed seedlings germinated *in vitro*, as commonly done in grow-out tests for seed sanitary analyses (Gitaitis and Walcott 2007). The isolation from seedlings permits to isolate a larger diversity of seed-borne microorganisms that are rare or in a viable but non cultivable state in the seed habitat. For microbial isolation only, seedlings germinated in sterile conditions were also obtained in a climatic chamber (25°C for tomato and bean and 20°C for wheat and rapeseed). The isolation was performed using 30 bean seedlings (7 days old), 15 rapeseed seedlings (4 days old), 6 tomato seedlings (7 days old) and 6 wheat seedlings (7 days old) that were crushed in 2 mL of sterile water to obtain a homogeneous suspension. The resulting suspensions were serial-diluted and plated and TSA10.

### Bacterial and fungal strain isolation protocols

For bacterial isolation, 250 μL of seed soaking and seedling suspensions were serially diluted in sterile water and plated on TSA10 (3 plates for each seed batch and 3 plates for each individual seedling). The choice of TSA10 as an isolation medium was motivated by previous culturomics studies in our team comparing multiple media and showing that this one recovered the largest seed bacterial diversity (Chesneau *et al*. 2022). The plates were incubated at 20°C for 7 days. Individual colonies showing differences in morphology were collected to maximize the bacterial diversity isolated for each sample. The number of isolates collected for each sample was: for tomato: 12 isolates from seed and 11 from seedling, for wheat: 10 from seed and 10 from seedling, for rapeseed: 46 from seed and 46 from seedling and for bean: 47 from seed and 46 from seedling. Each colony was transferred to a microplate containing 200 µL of tryptic soy broth (1 colony per well) and grown at 20°C for 3 days under agitation (70 rpm). Two negative culture controls, i.e. a well without inoculation, were included in each microplate. The cultures obtained were then preserved in glycerol 40% at -80°C and the taxonomic affiliation was performed after boiling the remaining bacterial suspensions (diluted 1/10 in sterile water, incubated 5 min at 99°C, then stored at -20°C). The boiled suspensions were used for library preparations of *gyrB* amplicon sequencing to identify the taxonomic affiliations and purity of the isolates (see below). Microbial isolation was performed on only 4 rapeseed genotypes (Aviso, Jupiter, Milena, Mohican) because we had an existing collection in the lab on the other six genotypes.

Filamentous fungi and yeast strain isolation have been performed on all plant genotypes and production conditions. The seed and seedling suspensions obtained were filtered using filtration units with mesh woven filters 31 μm (SEFAR NITEX 03-31/24). The supernatant containing the microorganisms that went through the filters (mainly bacteria) was discarded to isolate only larger organisms present on the filters (yeast and filamentous fungi). The filter was washed in 5 ml of sterilized osmotic water. The suspension was diluted in liquid malt medium 1.5% with tetracycline 10 mg/L (Sigma-Aldrich) and streptomycin 50 mg/L (Sigma-Aldrich) to prevent bacterial growth and dilutions (1/10, 1/50, 1/100, 1/200 and 1/500) were done in 96 - well plates. Plates were then incubated at 23°C for one week.

The following steps of the isolation procedure focused on microplates presenting microbial growth in half or less than half of the wells to maximize the chance to have pure culture in each well (Zhang J *et al*. 2021). When a well presented growth, the microbial suspension was collected using a micropipette and deposited on a 1.5% malt agar plate. Plates were incubated at 23°C until the observation of fungal development. For non-filamentous isolates that exhibited colony growth, their morphology was examined under a bright-field microscope to determine whether they were yeasts or bacteria resistant to the used antibiotics. A minimum of 20 isolates per treatment were selected to cover a large diversity of colony morphology. All the strains were stored at - 80°C in glycerol 30%.

### Seed microbiota characterization using amplicon sequencing

The bacterial and fungal communities of the seed samples were characterized using amplicon sequencing of the *gyrB* gene (bacteria) and ITS1 region (fungi). The bean samples were sequenced in one run and the tomato, rapeseed and wheat samples were sequenced together on a separate run.

DNA was extracted using the NucleoSpin® 96 Food kit (Macherey-Nagel, Düren, Germany) following the manufacturer’s instructions. The first PCR was performed with the primers gyrB_aF64/gyrB_aR353 (Barret *et al*. 2015), which target a portion of bacterial *gyrB* gene, and ITS1F/ITS2 primers (Buée *et al*. 2009), which target fungal ITS1 region. PCR reactions were performed with a high-fidelity Taq DNA polymerase (AccuPrimeTM Taq DNA polymerase Polymerase System, Invitrogen, Carlsbad, CA, USA) using 5 µL of 10X Buffer II, 1 µL of forward and reverse primers (100 µM for *gyrB* and 10 µM for ITS1), 0.2 µL of Taq and 5-10 µl of DNA. For gyrB_aF64/gyrB_aR353 primers, PCR cycling conditions were done with an initial denaturation step at 94°C for 3 min, followed by 35 cycles of amplification at 94°C (30 s), 55°C (45 s) and 68°C (90 s), and a final elongation at 68°C for 10 min. For ITS1F/ITS2 primers, the cycling conditions were an initial denaturation at 94°C for 3 min, followed by 35 cycles of amplification at 94°C (30 s), 50°C (45 s), and 68°C (90 s), and a final elongation step at 68°C for 10 min. Amplicons were purified with magnetic beads (Sera-MagTM, Cytiva, Marlborough, MA, USA). The second PCR was conducted to incorporate Illumina adapters and barcodes. The PCR cycling conditions were the same for the two markers: denaturation at 94°C (1 min), 12 cycles at 94°C (1 min), 55°C (1 min) and 68°C (1 min), and a final elongation at 68°C for 10 min. Amplicons were purified with magnetic beads, quantified with QuantIT PicoGreen dsDNA Assay Kit (Invitrogen) and pooled in equimolar concentration. Pool concentration was measured with quantitative PCR (KAPA Library Quantification Kit, Roche, Basel, Switzerland). Amplicon libraries were mixed with 5-15% PhiX and sequenced with two MiSeq reagent kits v3 600 cycles (Illumina, San Diego, CA, USA) for 2 x 250 bp on the Illumina MiSeq platform (ANAN platform, SFR QuaSav, Angers, France). A blank extraction kit control, a PCR-negative control and PCR-positive control (*Lactococcus piscium* DSM6634, a fish pathogen that is not plant-associated) were included in each PCR plate. The raw amplicon sequencing of the seed microbiota data are available on the European Nucleotide Archive (ENA) with the accession number PRJEB59579 for bean samples and PRJEB91550 for rapeseed, tomato and wheat samples.

### Isolate sequencing for taxonomic identification

Bacterial identification was done using *gyrB* amplicon sequencing as described above for the seed samples. The protocol was adapted to enable the sequencing of 1500 bacterial isolates on the same run. A plate number index was added to the reverse primer of the gyrB_aF64/gyrB_aR353 primer set of the first PCR to increase multiplexing capacity. A total of 32 plates each containing 92 or 93 isolates were characterized, distributed as follows: 16 plates for bean, 4 plates for tomato, 8 plates for rapeseed and 4 plates for wheat. The bean isolates were sequenced in one run and the tomato, rapeseed and wheat isolates were sequenced together on a separate run.

PCR reactions were performed with GoTaq® G2 DNA Polymerase (Promega, Madison, WI, USA) using 5 µL of 5X Reaction Buffer, 0,5 µL of each forward and reverse primers (50 µM), 0.5 µL of dNTP mix (10 mM), 0.125 µL of Taq and 2.5 µL of bacterial boiled suspensions. PCR cycling conditions were done with an initial denaturation step at 94°C for 3 min, followed by 25 cycles of amplification at 94°C (30 s), 55°C (45 s) and 72°C (90 s), and a final elongation at 72°C for 10 min. Amplicons were purified with magnetic beads (Sera-MagTM, Cytiva, Marlborough, MA, USA). The second PCR was conducted to incorporate Illumina adapters and barcodes. The PCR cycling conditions were: denaturation at 94°C (1 min), 12 cycles at 94°C (1 min), 55°C (1 min) and 72°C (1 min), and a final elongation at 72°C for 10 min. Amplicons were purified with magnetic beads and pooled. Pool concentration was measured with quantitative PCR (KAPA Library Quantification Kit, Roche, Basel, Switzerland). For bean run, amplicon libraries were mixed with 10% PhiX and sequenced with MiSeq reagent kit v2 Nano 500 cycles (Illumina). For tomato, rapeseed and wheat run, amplicon libraries were mixed with 15% PhiX and sequenced with MiSeq reagent kit v2 500 cycles (Illumina). A PCR-negative control and PCR-positive control (*Lactococcus piscium* DSM6634, a fish pathogen that is not plant-associated) were included in each PCR plate.

Filamentous fungi identification was based on the internal transcribed spacer (ITS) region using primers NS7 (5’-GAGGCAATAACAGGTCTGTGATGC-3’) and ITS4 (5’-TCCTCCGCTTATTGATATG-3’) (White *et al*. 1990). For Basidiomycota yeasts, the identification was enhanced using the sequence of the D1/D2 region of the large 26S rDNA subunit using primers LROR (5’-ACCCGCTGAACTTAAGC-3’) and LR5 (5’-TCCTGAGGGAAACTTCG-3’) (Vilgalys and Hester 1990). Gene sequences were amplified by colony PCR using GoTaq® G2 Hot Start Polymerase from Promega Corporation in concordance with manufacturer’s instructions and Sanger sequenced using Eurofins Genomics facilities.

### Bioinformatics, statistics and diversity analyses

#### Processing for seed microbiota data

Each sequencing run was independently processed with the standardized bioinformatic pipeline used for the Seed Microbiota Database (Simonin *et al*. 2022) using QIIME 2 and DADA2 (Callahan *et al*. 2016, Bolyen *et al*. 2019). In brief, primer sequences were removed with CUTADAPT 2.7 and trimmed FASTQ files were processed with DADA2 v.1.10, using variable truncation parameters depending on the sequencing quality of the run. Chimeric sequences were identified and removed with the removeBimeraDenovo function of DADA2. ITS1 were processed using only the forward reads as recommended by Pauvert *et al*. (2019) to maximize fungal diversity detection. Amplicon sequence variant (ASV) taxonomic affiliations were performed with a scikit-learn multinomial naive Bayesian classifier implemented in QIIME 2 (qiime feature-classifier classify-sklearn; Bokulich *et al*. 2018). *gyrB* reads were classified with an in-house *gyrB* database (train_set_gyrB_v5.fa.gz) available upon request. The ASVs derived from ITS1 reads were classified with the UNITE v.8 fungal database (Nilsson *et al*. 2019). Nonfungal ASVs were filtered from the ITS gene studies. Since the primer set gyrB_aF64/gyrB_aR353 primers can sometimes coamplify *parE*, a paralog of *gyrB*, the *gyrB* taxonomic training data also contained *parE* sequences. ASVs affiliated with *parE* or unclassified at the phylum-level were filtered from the dataset. ASVs with less than 20 reads and detected in only one sample were removed from the dataset. After these filtering steps, samples with < 1000 reads were excluded from the dataset.

Diversity and community structure analyses were performed in R 4.3.1 using the Phyloseq (v.1.44.0), Vegan (v.2.6-6.1) and Microbiome (v.1.22) packages (Oksanen *et al*. 2007, McMurdie and Holmes 2013, Lahti, Shetty, and Blake 2017). For the alpha diversity analysis, the datasets were rarefied to the lowest number of reads observed in a sample: 5177 reads for gyrB and 9981 reads for ITS1. At these rarefaction levels, the sequencing depth was sufficient to reach a plateau on the rarefaction curves Seed microbiota diversity was explored using ASV richness. Changes in seed microbiota structure were assessed on log_10_+1 transformed values using Bray–Curtis dissimilarity and principal coordinate analysis (PCoA) was used to plot the ordination. We assessed statistical effects of plant genotype and production mode for each crop independently on the ASV richness and bacterial community size using generalized linear models (*glm* and *glm*.*nb* functions in lme4 package v.1.1-35.3) and *post hoc* comparisons were performed using the Tukey method (package emmeans v.1.10.2) in R. The effects on seed microbiota structure were analyzed using a PERMANOVA with the *adonis2* function in Vegan. Sequences were aligned with DECIPHER 2.28.0 (Wright 2016), and neighbor-joining phylogenetic trees were constructed with phangorn 2.11.1 (Schliep 2011) for visualization of the seed microbiota diversity and the position of strains isolated on these phylogenies.

#### Data processing for bacterial and fungal strain collection taxonomic identification

For bacterial isolate identification using *gyrB* gene amplicon sequencing, the raw reads were demultiplexed using cutadapt (-e 0 -O 20) with plate number index. The reads denoising was performed with DADA2 v.1.28.0 in R with the following parameters for the filterAndTrim function (truncLen:190, 130, maxEE=2, truncQ=2, maxN=0). Chimeric sequences were identified and removed with the removeBimeraDenovo (method=“consensus”) function of DADA2. *gyrB* reads were classified with an in-house *gyrB* database (train_set_gyrB_v5.fa.gz). A custom made script was made to identify in each sample the three most abundant unique sequences (Amplicon Sequence variants, ASVs) and their taxonomy. A conservative approach was used to select bacterial isolates included in the final culture collection based on taxonomic affiliation results, following these criteria: isolates were selected if they had a minimum of 60 reads in sample, the most abundant ASV represented > 90% of reads, the taxonomic affiliation was similar between the most abundant and second most abundant ASV (due to *parE* paralog of *gyrB* or sequencing errors). Based on these criteria, 84% of the isolates were included in the final bacterial collection presented here (2510 validated isolates on 2952 sequenced).

Taxonomic affiliation of fungal isolates at the genus level were obtained using DADA2 and the UNITE v.8 fungal database (Nilsson et al., 2019). The refined species identification of the Basidiomycota’s yeast isolates methodology is described in detail in Marchi et al., 2026.

## RESULTS

### 1. Seed Microbiota Characterization

#### a. Contrasted bacterial community sizes between plant species, genotypes and production modes

First, we characterized the cultivable bacterial community sizes of the different seed samples by plating seed macerates on solid medium and counting CFUs (Figure 1A). Seed samples harbored very contrasted levels of bacterial colonization ranging from 10 to 100 million CFUs per gram of seeds. Tomato, rapeseed and wheat had similar ranges of CFU values (average log 5.29, 5.77 and 5.47, respectively), and bean seeds presented lower community sizes (average log 3.85).

**Figure 1:**
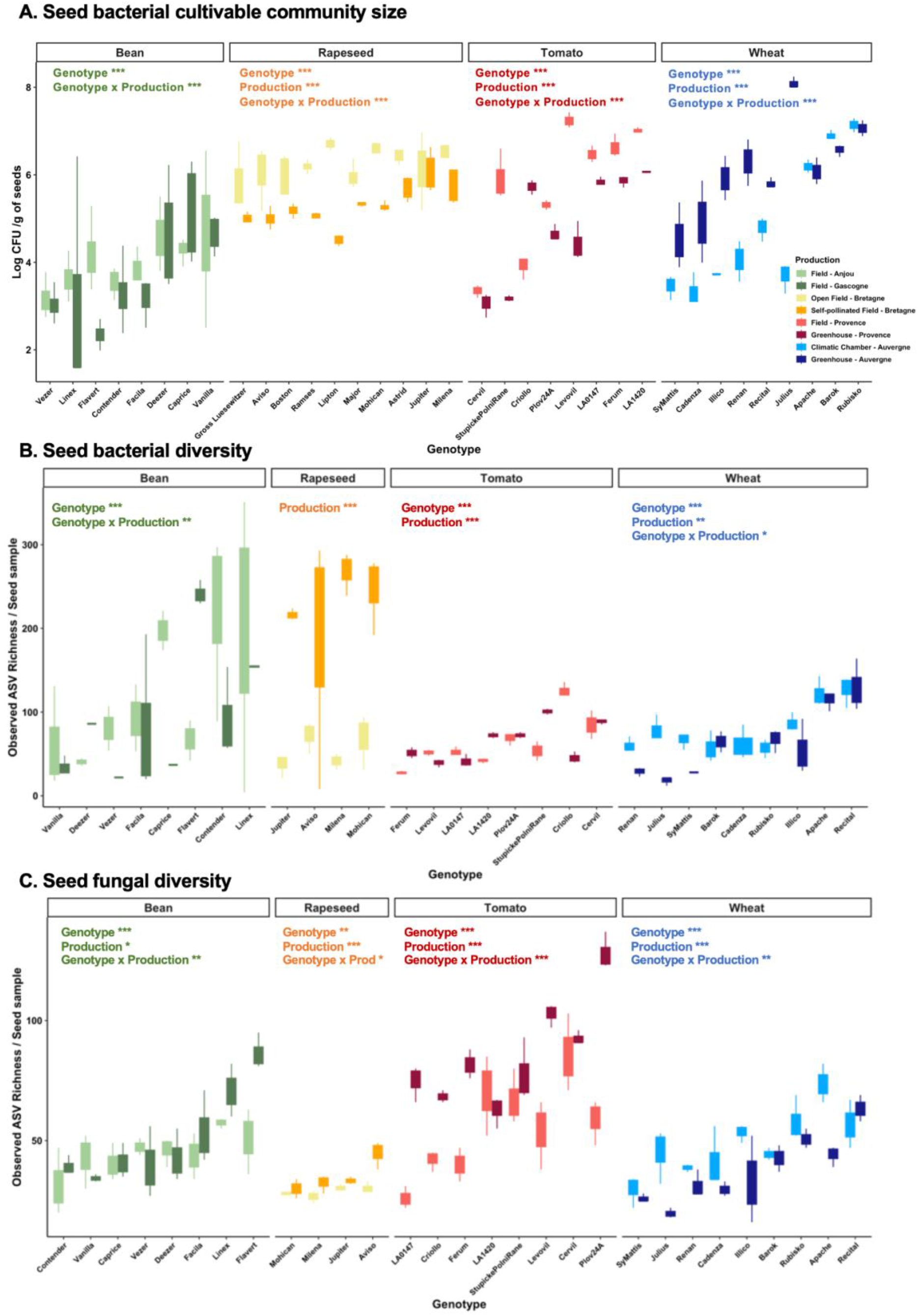
A. Bacterial community size of seed samples assessed by plating seed macerates on 10% strength TSA medium (CFU: Colony Forming Unit). B. Seed bacterial richness estimated with *gyrB* Amplicon Sequence Variants (ASVs). C. Seed fungal richness estimated with ITS1 ASVs. Statistical analyses have been done using *glm* for each plant species separately and the significant variables are indicated on top of each panel by asterisks (* < 0.05; ** < 0.01; *** < 0.001).

We assessed the influence of the genotype and production mode on seed bacterial community sizes for each plant species independently, since the four species were produced following specific culture conditions and in different localizations. For bean, the genotype and the interaction between genotype and production were significant (P<0.001), but not the production mode alone (P=0.10). These effects were mostly driven by a gradient of bacterial population sizes from Vezer (lowest values) to Vanilla genotype (highest values), and lower CFUs in Gascogne for three genotypes (Flavert, Linex, Deezer). For rapeseed, all variables and their interaction were significant drivers of bacterial community size, but the production mode had the largest effect (P<0.001). Overall, seeds produced in open field conditions had a higher bacterial colonization than the self-pollinated seeds. For tomato, very large differences in bacterial CFUs were observed between genotypes and production conditions (ranging from log 2 to 7, P<0.001 for all factors and their interaction). For all genotypes except Cervil, seeds produced in the field had higher bacterial colonization than greenhouse-produced seeds (1 to 3 logs of difference). For wheat, all variables and their interaction were significant drivers of bacterial community size, with both production mode and genotype having large effects (P<0.001). For six out of nine genotypes, seeds produced in the greenhouse had higher bacterial colonization than in the climatic chamber production. Altogether, these results show that seed bacterial colonization is strongly influenced by both genotype and production mode for the four crop species considered.

#### b. Seed bacterial and fungal diversities are highly influenced by host genotypes and production modes

Next, we characterized the seed microbiota diversity of the different seed samples using metabarcoding. The samples presented correspond to the same that were used next for isolation of bacterial and fungal strains (see below). Hence, only four rapeseed genotypes are presented here (the ones on which isolation was done) and richness results are expressed as observed ASVs per seed sample (and not per g of seeds) to be directly comparable to the strain collection results presented in the next section.

For bacterial communities, the ASV richness ranged from 4 to 351 ASVs per seed sample (Figure 1B), with an average of 89.5 ASVs and a total of 3145 ASVs detected in the dataset across all seed samples (3099 ASVs after rarefaction). Tomato seeds had the lowest richness values on average (56.5 ASVs), compared to wheat (71 ASVs), bean (87.5 ASVs) and rapeseed (90 ASVs). Bean and rapeseed seed samples had extremely variable bacterial richness, even between replicates of the same condition, while tomato and wheat had more homogeneous richness values.

Bean bacterial richness was significantly influenced by genotype (P<0.001) and its interaction with production mode (P=0.008). For rapeseed, only production mode influenced bacterial richness (P<0.001) with a higher ASV richness observed in the self-pollinated field for the four genotypes (treatment using fly pollination in insect proof cages to avoid cross-pollination between genotypes). For tomato seeds, bacterial richness was structured by the genotype (P<0.001) and its interaction with production mode (P<0.001). For wheat, genotype, production modes and their interaction were significantly modulating bacterial richness (P<0.001).

For seed fungal communities, the ASV richness ranged from 16 to 138 ASVs per seed sample (Figure 1C), with an average of 50.8 ASVs and a total of 881 ASVs detected in the dataset (876 ASVs after rarefaction). The variability between samples was less pronounced for fungal richness than bacterial communities and the four species had comparable richness ranges. Rapeseed seeds had the lowest richness values on average (30 ASVs), compared to wheat (43.5 ASVs), bean (46.5 ASVs) and tomato (68.5 ASVs). For the four plant species, all factors and their interaction were significant (P<0.001).

For bean fungal richness, the genotype effect was mainly driven by significant differences between Vanilla, Facila, Linex and Flavert in the Gascogne field. For rapeseed, production was the most influential factor (P<0.001) with a higher ASV richness observed in the self-pollinated field. For tomato seeds, production mode and the interaction with genotype were the strongest drivers, with a higher fungal richness observed in the greenhouse than in the field for five out of eight genotypes. For wheat, similar to bacterial communities, genotypes harbored a gradient of richness with Apache and Recital having the highest diversity. The significant interaction between genotype and production was due to three genotypes exhibiting higher fungal richness in the climatic chamber compared to the greenhouse production (Julius, Cadenza, Illico, Apache).

Next, we found that bacterial and fungal richness values were not correlated across all samples (Figure S2, Spearman test: R^2^=0.02 P=0.09). Most seed samples had a higher bacterial richness (rapeseed, wheat, bean) than fungal richness, with the exception of tomato seeds which harbored more fungal diversity especially in the greenhouse production.

#### c. Distinct seed bacterial and fungal community structures and compositions depending on plant species and production

Seed microbiota composition was next assessed for the four plant species (Figure 2 and 3). Distinct bacterial community composition between the different plant species were observed (Figure 2A, R^2^=0.29, P<0.001), especially for tomato seeds which clustered distantly from the other three crops along axis 1. For bean, tomato and wheat, the genotype and its interaction with production mode were the main drivers of community structure (Figure 2B-2D-2E, R^2^=17 to 38%, P<0.001) and the production alone was a secondary driver (R^2^=5 to 11%). In contrast, the rapeseed bacterial community was mainly structured by the production mode (Figure 2C, R^2^=28%) and secondly by the interaction between genotype and production (R^2^=14%).

**Figure 2:**
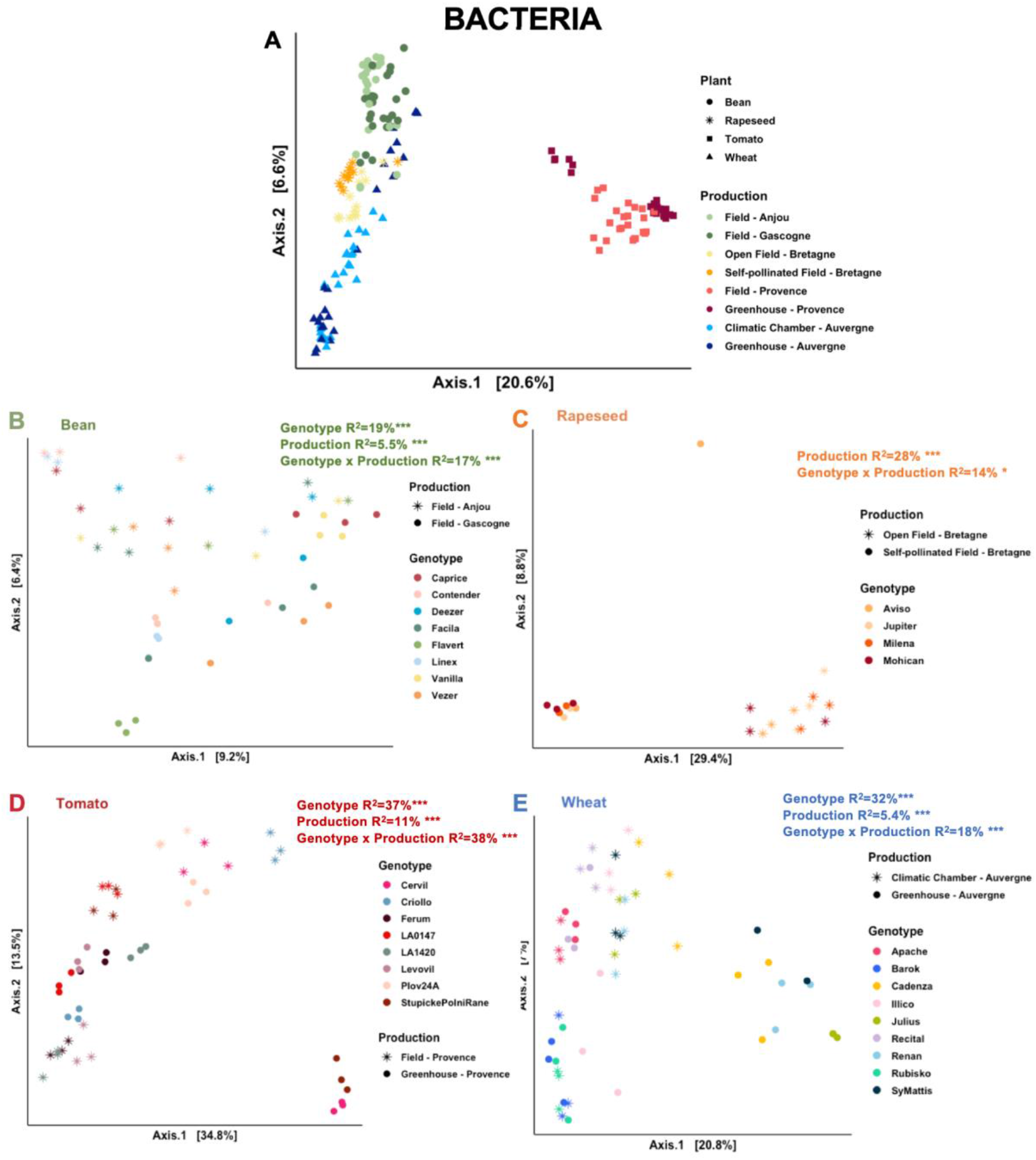
Seed bacterial community structure of the four plant species. A. Global ordination (PCoA) based on Bray-Curtis distances of all seed samples. The following panels correspond to an independent ordination per plant species: B. bean, C. rapeseed, D. tomato, E. wheat. The effects of genotype and production modes were assessed using a Permanova and the results for each plant species are indicated on the top right corner of the panel.

**Figure 3:**
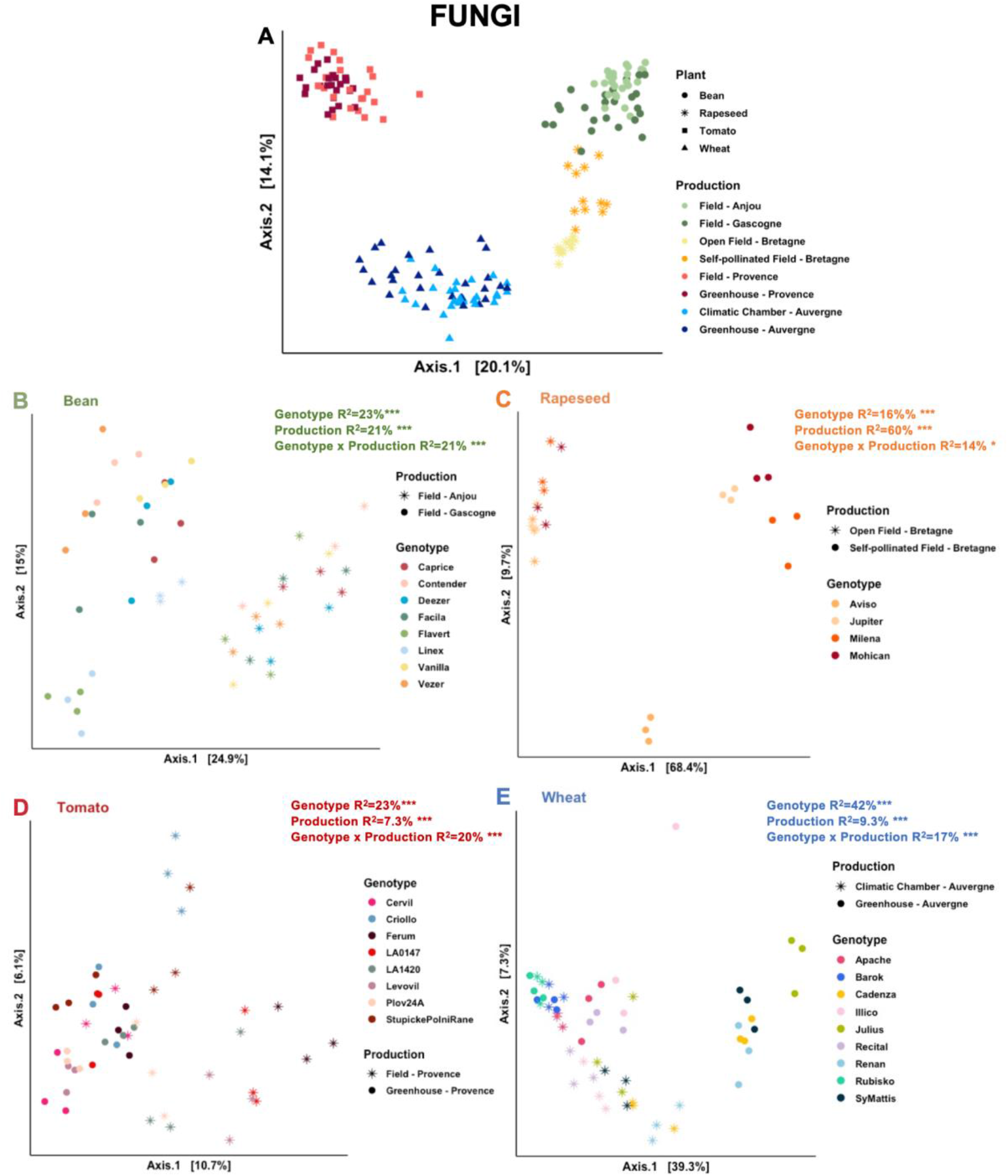
Seed fungal community structure of the four plant species. Similar legend description as Figure 2.

Distinct fungal community compositions were also observed between the different plant species (Figure 3A, R^2^=0.39, P<0.001). For the four plant species, all factors and their interaction were significant but the variance explained and their hierarchy of factors varied between species (Figure 3B through 3D).

For wheat and tomato, the genotype and its interaction with production mode were the main drivers of community structure (Figure 3D-3E, R^2^=17 to 42%, P<0.001) and the production alone was a secondary driver (R^2^=7 to 9%). Again, the rapeseed bacterial community was highly structured by the production mode (R^2^=60%) and secondly by the genotype (R^2^=16%) and the interaction between genotype and production (R^2^=14%) (Figure 3C). Bean fungal community was equally influenced by genotype (R^2^=23%), production (R^2^=21%) and their interaction (R^2^=21%) and production alone (Figure 3B).

Next, we analyzed the taxonomic compositions of the bacterial (Figure 4 A-D) and fungal communities (Figure 5 A-D) at the genus level. All plant species had clearly distinct taxonomic profiles, even if some ASVs were shared between the different plant species (Figure 4E-F, 5E-F).

**Figure 4:**
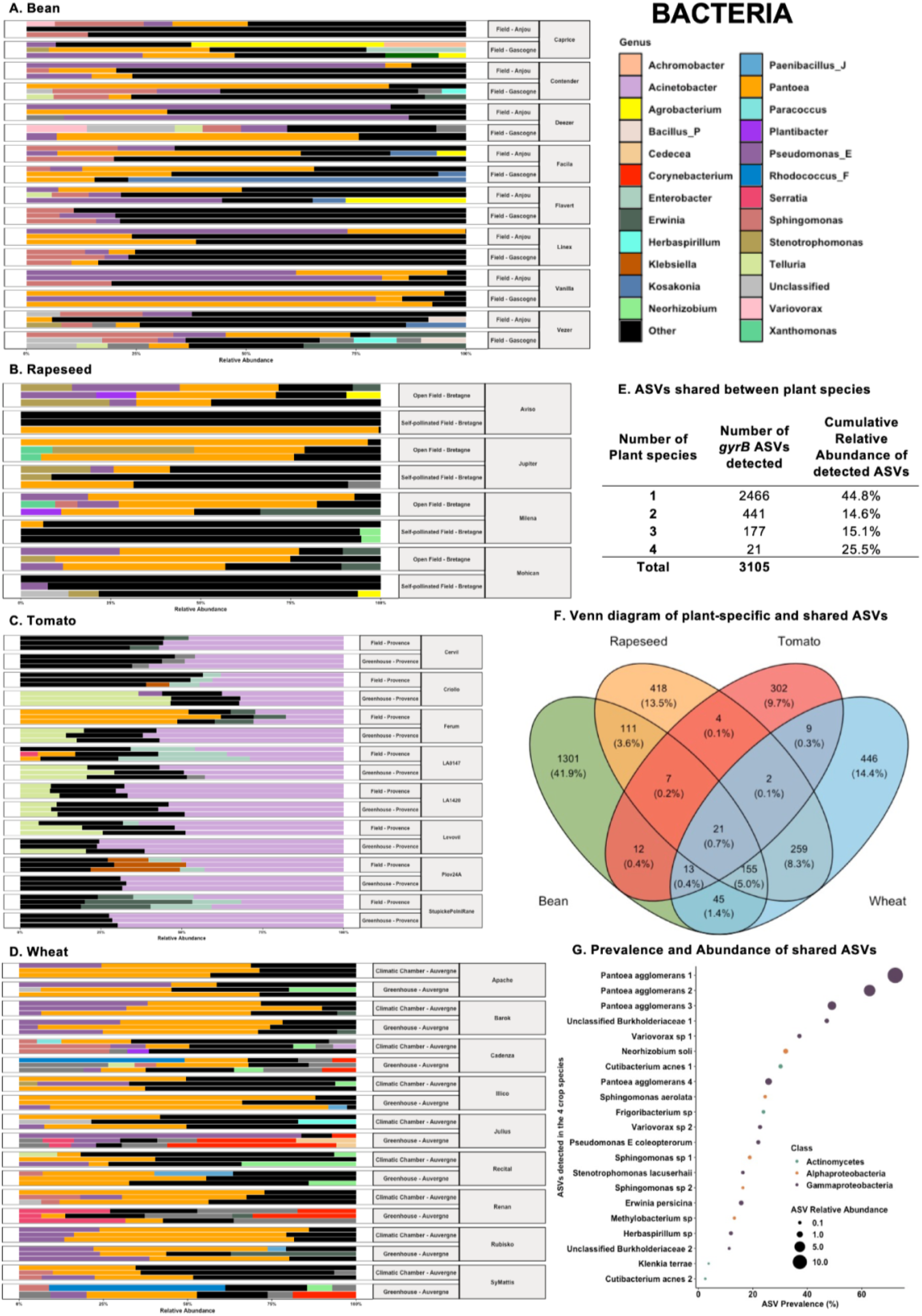
A-D) Bacterial taxonomic profiles by plant species (A: bean, B: rapeseed, C: Tomato, D: Wheat). Capital letters (ex: _P) indicate the genus taxonomic group. E) Number and cumulative relative abundance of ASVs specific to one plant species or shared with 2, 3, or 4 plant species. F) Venn diagram representing the number of ASVs specific or shared between species The values in parenthesis indicate the percentage of the bacterial diversity found in each intersection. G) Prevalence and cumulative relative abundance across all samples for the 21 ASVs shared between the 4 plant species.

**Figure 5:**
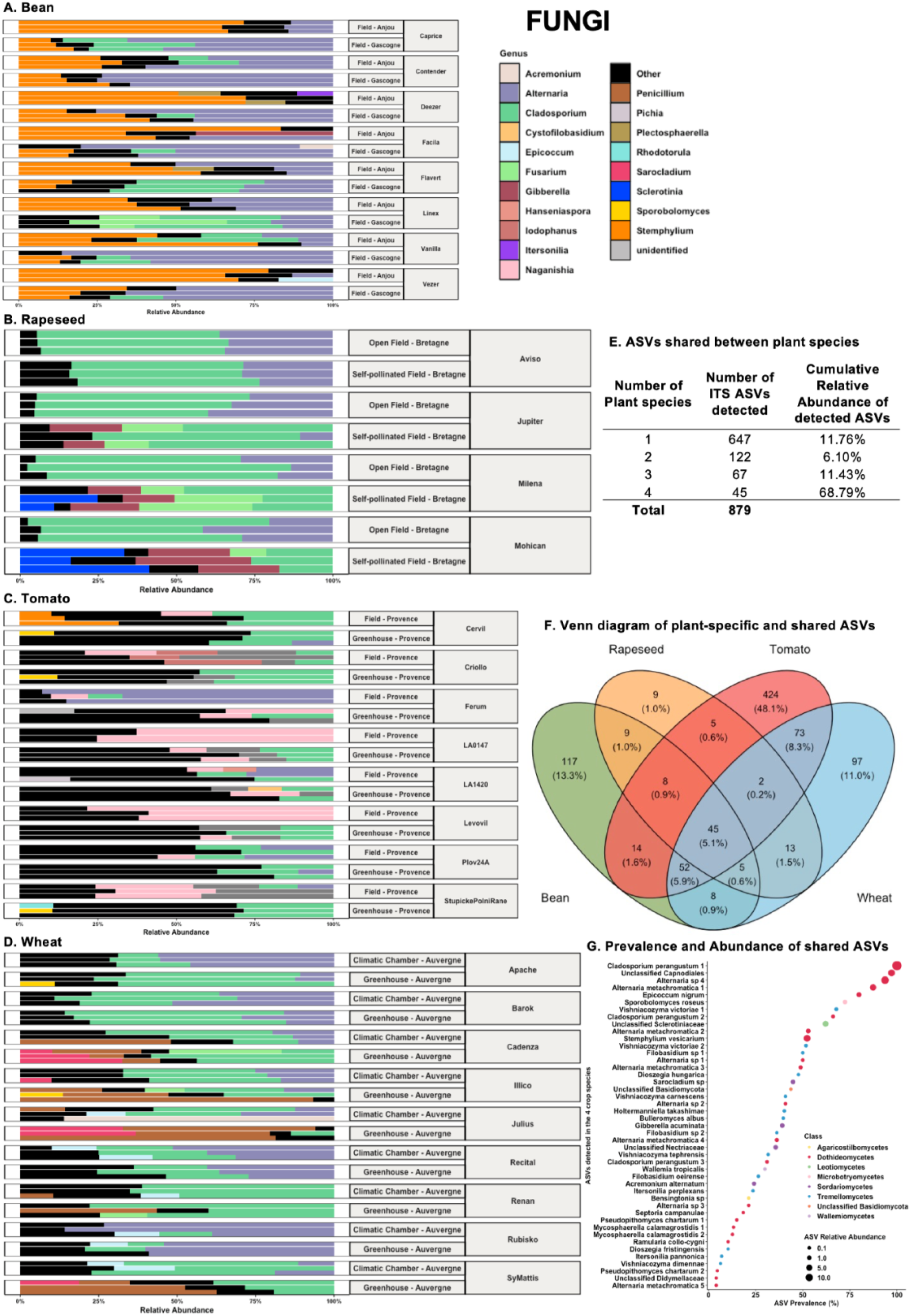
A-D) Bacterial taxonomic profiles by plant species (A: bean, B: rapeseed, C: Tomato, D: Wheat). E) Number and cumulative relative abundance of ASVs specific to one plant species or shared with 2, 3, or 4 plant species. F) Venn diagram representing the number of ASVs specific or shared between species. G) Prevalence and cumulative relative abundance across all samples for the 21 ASVs shared between the 4 plant species.

Bean bacterial community compositions were highly variable between conditions and also between replicates, but the *Pantoea, Sphingomonas* and *Pseudomonas* (E group) genera were the most abundant (Figure 4A). Rapeseed communities were less variable and presented clear signatures according to production mode (Figure 4B). In open field samples, *Pantoea* and *Pseudomonas* were dominant, while no clear signature was visible in self-pollinated samples due to the higher diversity and variability. Tomato bacterial communities were consistently dominated by *Acinetobacter* across all samples (Figure 4C). Under specific genotype or production conditions, the *Telluria, Pantoea, Erwinia, Klebsiella* and *Enterobacter* genera were also abundant. Finally, wheat bacterial communities were generally dominated by *Pantoea*, associated with a variable abundance of *Pseudomonas, Sphingomonas, Corynebacterium* and *Neorhizobium* depending on the conditions (Figure 4D).

When analyzing the presence-absence of each bacterial ASV across the different plant species, we found that the majority of ASVs were detected in one plant species (n=2466 on 3105) representing 44.8% of the relative abundance of the dataset (Figure 4E). A large fraction of these plant specific ASVs were associated with bean (1301 ASVs, Figure 4F) which harbor highly variable and high richness communities. The remaining ASVs (n=639) were shared between 2 to 4 species, especially between wheat, rapeseed and bean (n=570), while tomato samples shared few ASVs. Interestingly, only 21 ASVs belonging to three classes (*Actinomycetes, Alphaproteobacteria, Gammaproteobacteria*) were detected in the four plant species but the cumulative abundance of these shared ASVs was significant (25.5%). The prevalence and relative abundance across all samples of these 21 shared ASVs is presented in Figure 4G. The 3 most prevalent ASVs were affiliated to the *Pantoea agglomerans* species and the other shared ASVs were affiliated to a dozen of species, including *Variovorax sp, Neorhizobium sp, Cutibacterium sp* or *Sphingomonas sp*.

Bean fungal community compositions were less variable than bacterial profiles, with a clear dominance of *Alternaria, Stemphylium* and *Cladosporium* (Figure 5A). Rapeseed communities presented again clear signatures according to production mode but with a high abundance of *Cladosporium* in all samples (Figure 5B). *Gibberella* and *Sclerotinia* genera were specifically abundant in self-pollinated samples. Tomato fungal communities were highly variable but *Cladosporium* was abundant in most conditions and *Naganishia* was often dominant under field conditions (Figure 5C). Finally, wheat fungal communities were generally dominated by *Cladosporium* and *Alternaria* (Figure 5D), under greenhouse conditions *Penicillium* and Sarocladium were abundant for specific genotypes (Cadenza, Julius, SyMattis).

The patterns of presence-absence of fungal ASV differed from bacterial ones across the 4 plant species. While the majority of ASVs were detected in one plant species (n=647 on 879), they only represented 11.76% of the relative abundance of the dataset (Figure 5E). These plant specific ASVs were mainly associated with tomato (424 ASVs, Figure 5F) which harbor highly variable and diverse fungal communities The fungal ASVs contributing the largest relative abundance (68.79%) were the 45 ASVs shared between the 4 plant species. The 45 shared ASVs included extremely prevalent taxa (> 70% of samples, Figure 5G), such as filamentous fungi *Cladosporium perangustum, Alternaria* (*chromatica &* sp), *Epicoccum nigrum* or Basidiomycota yeast *Sporobolomyces roseus* and *Vishniacozyma victoriae*.

Altogether, these results show that seed bacterial and fungal communities were highly contrasted between the genotypes and production modes for the four crop species considered. These contrasted seed samples were thus a valuable resource to obtain a diversified seed microbial strain collection.

### Part 2: Seed-borne strains isolated from the four plant species

#### Bacterial strain collection

A total of 2510 bacterial isolates were identified and validated as pure strains to constitute the SUCSEED bacterial collection. The strains isolated from bean were the most numerous (Figure 6A, n=1276, representing 616 unique gyrB ASVs), followed by rapeseed (n=636, 326 unique ASVs), tomato (n=292, 143 unique ASVs) and wheat (n=292, 183 unique ASVs). These numbers are consistent with the sampling efforts (i.e number of colonies picked per sample) and bacterial richness observed (Figure 1B), which were higher for bean and rapeseed. The total number of unique strains ASVs isolated per condition (i.e a genotype for a given production mode) based on isolates recovered from both seeds and seedlings is presented in Figure 6B. For bean and rapeseed collections, we isolated 55.9-54.9 unique ASVs on average and 16.5-15.3 for tomato and wheat per condition, respectively. This culturing effort was able to isolate 10% (wheat) to 21% (tomato) of the seed microbiota diversity detected by metabarcoding (Figure 6C). While these values can appear low, this incomplete recovery is mainly driven by the difficulty to isolate rare taxa. The most prevalent and abundant bacterial taxa (Top 50) were successfully isolated and were recovered in high proportions for all plant species (46 to 72%, Figure 6C). Collectively, the isolated bacterial ASVs represented 69 to 76% of the seed microbiota relative abundance assessed in metabarcoding, indicating that the bacterial strain collection recovered a large fraction of the seed bacterial composition. Interestingly, we also isolated a large number of strains for which the ASVs were not detected in metabarcoding (Figure 6D). Indeed, 44 to 60% of our isolates were not detected in metabarcoding despite the good sequencing depth obtained in the initial characterization of seed microbiota (Figure S1). This observation suggests that culturomics offer a complementary characterization of seed microbiota permitting the detection of rare taxa or having low amplification in PCR.

**Figure 6:**
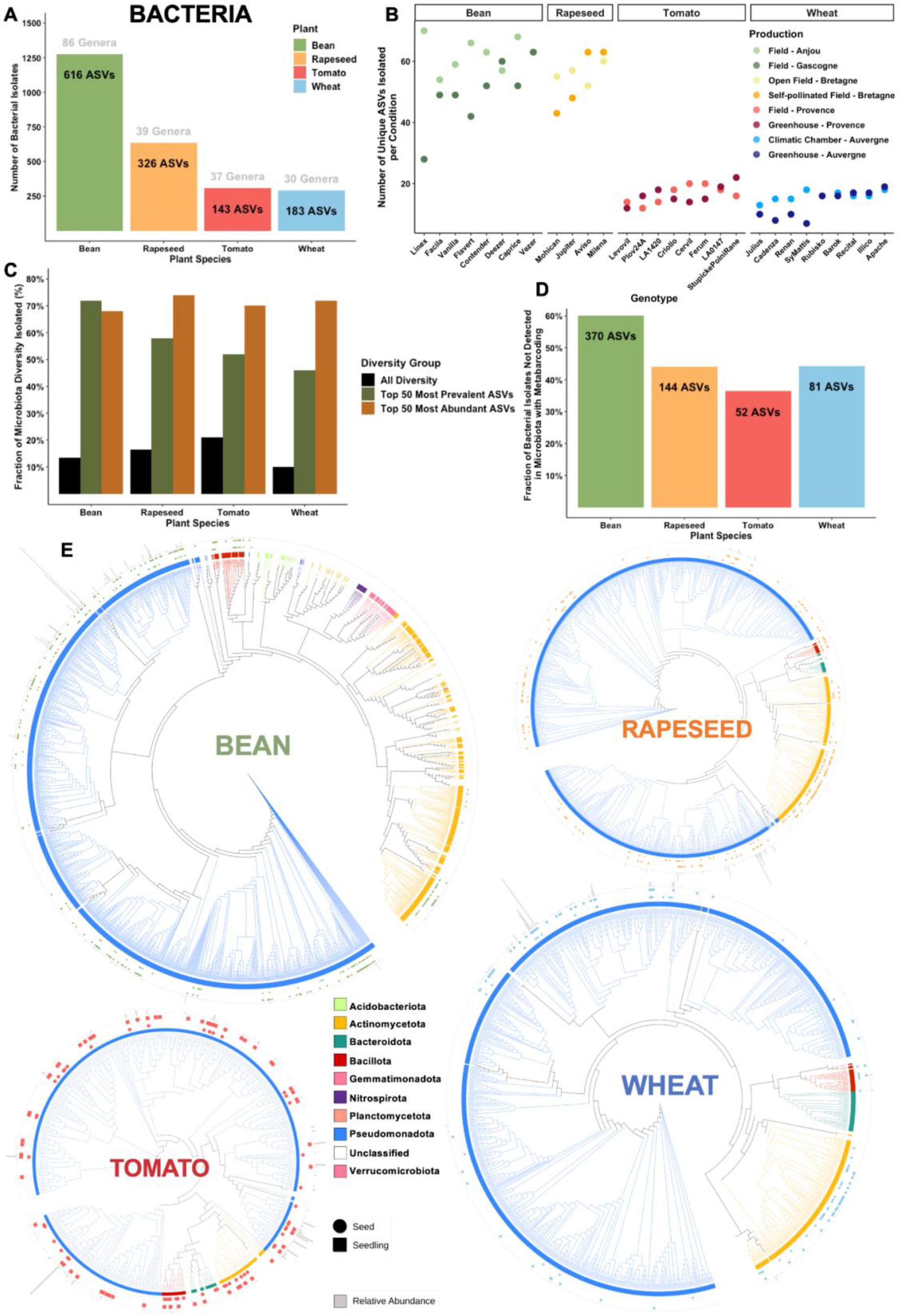
A) Total number of bacterial strains isolated by plant species. The number of unique ASVs and genera is indicated for each plant species. B) Number of unique strains ASVs isolated for each condition (i.e a genotype in a given production mode) based on both seed and seedling isolates. C. Fraction of seed bacterial diversity successfully isolated, based on the comparison of strains ASVs and all ASVs detected in metabarcoding. D) Fraction of strains ASVs only found in the strain collection and not in the metabarcoding analysis. E) Phylogenetic trees of all *gyrB* ASVs detected in seed microbiota based on metabarcoding analysis for each plant species separately. ASVs for which a bacterial isolate is available in the collection are indicated by a symbol (circle for seed isolates and square for seedling isolates). The colored inner circle represents the ASV taxonomic affiliation at the phylum level and vertical grey bars on the outer circle represent the average relative abundance of each ASV.

Next, we assessed the phylogenetic representativeness of the strains isolated compared to seed microbiota diversity characterized using metabarcoding (Figure 6E) by representing all the *gyrB* ASVs detected on a phylogenetic tree for each plant species. The symbols (square and circle) represent when the ASVs have been successfully isolated from seeds or seedlings and the grey bars their relative abundance detected in microbiota by metabarcoding. The majority of isolates recovered belonged to *Pseudomonadota* which is dominant for all plant species, followed by *Actinomycetota* and *Bacillota*. The other phyla (e.g *Bacteroidota, Acidobacteriota, Gemmatimonadota*) had very low abundance and were not successfully recovered in our collection, with the exception of some *Bacteroidota* from tomato seeds. The visualization of the trees confirms the successful isolation of most ASVs with high relative abundance, often from both seeds and seedlings, indicating their ability to transmit to the next generation.

The number of isolates by bacterial genus obtained from the four plant species is presented in Figure 7. We highlight the number of isolates recovered from seeds and seedlings in different colors. While the culturing effort from seeds and seedlings was similar for all plants (Figure S3A), we clearly observed that some bacterial classes were more frequently isolated from either seeds or seedlings (Figure S3B). *Alpha*- and *Gammaproteobacteria* were more frequently isolated from seedlings (68% and 58% of isolates, respectively), but *Actinomycetes* were more commonly isolated from seeds (63% of isolates), suggesting different abundance and transmission patterns of these taxonomic groups.

**Figure 7:**
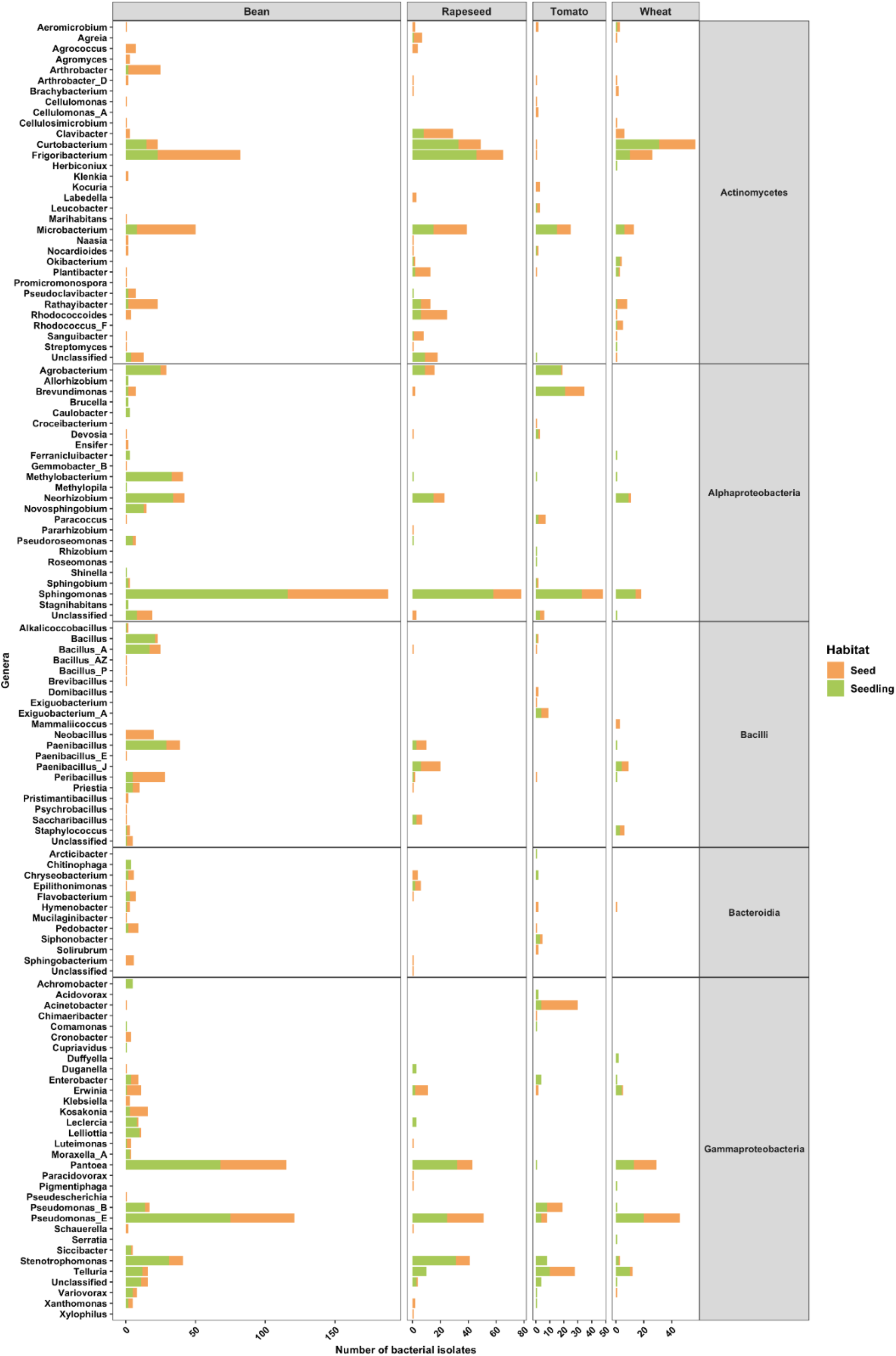
List of all bacterial genera isolated from the four plant species. For each genus the number of isolates colored by their source of isolation (seed or seedling germinated *in vitro*) is indicated. The genera are organized by bacterial classes.

Isolates affiliated to *Pseudomonas, Pantoea Sphingomonas, Microbacterium, Telluria* and *Stenotrophomonas* were recovered from all 4 plant species. Out of the 21 shared ASVs between the four species previously identified (Figure 4G), 16 of them were successfully isolated including the four *Pantoea agglomerans* taxa. For the other genera, their isolation varied more between plant species. For instance, *Curtobacterium, Frigoribacterium, Neorhizobium and Rathayibacter* were frequently isolated from bean, rapeseed and wheat, but not from tomato. On the opposite, *Acinetobacter* and *Brevundimonas* genera were more frequently isolated from tomato than the three other species. We obtained a high number of isolates of *Bacillus and Neobacillus* from bean seeds only or *Rhodococcoides* and *Clavibacter* mainly from rapeseed. These observations are in line with the metabarcoding results showing higher similarity between bean, rapeseed and wheat, while tomato is more distinct, but also that each species harbors specific taxa (Figure 2A, Figure 4).

#### Fungal strain collection

A total of 837 fungal isolates were identified and validated as pure strains to constitute the SUCSEED fungal collection. The strains isolated from tomato were the most numerous (Figure 8A, n=249, representing 58 unique ITS1 ASVs), followed by wheat (n=207, 37 unique ASVs), bean (n=193, 26 unique ASVs) and rapeseed (n=188, 19 unique ASVs). These numbers are consistent with the fungal richness observed (Figure 1C), which were higher for tomato and lower for rapeseed. The total number of unique strain ASVs isolated per condition (i.e a genotype for a given production mode) based on isolates recovered from both seeds and seedlings is presented in Figure 8B. For bean, rapeseed and wheat collections, we isolated 4 to 7 unique ASVs on average and 20 for tomato but this was driven by a higher number of ASVs recovered in the greenhouse production, especially the Plov24a genotype (42 ASVs). This culturing effort was able to isolate 6% (rapeseed) to 18% (tomato) of the seed microbiota diversity detected by metabarcoding (Figure 8C). These values are similar to the bacterial collection, and also reflect the difficulty to isolate rare taxa. A third of the most prevalent and abundant fungal taxa (Top 50) were isolated and were recovered in similar proportions for the different plant species (26 to 42%, Figure 8C). Collectively, the isolated fungal ASVs represented 74 to 82% of the seed microbiota relative abundance assessed in metabarcoding, indicating that the fungal strain collection recovered a significant fraction of the seed fungal composition. Compared to bacterial isolates, only few fungal isolates were not detected in metabarcoding, except for tomato seeds (34%, Figure 8D).

**Figure 8:**
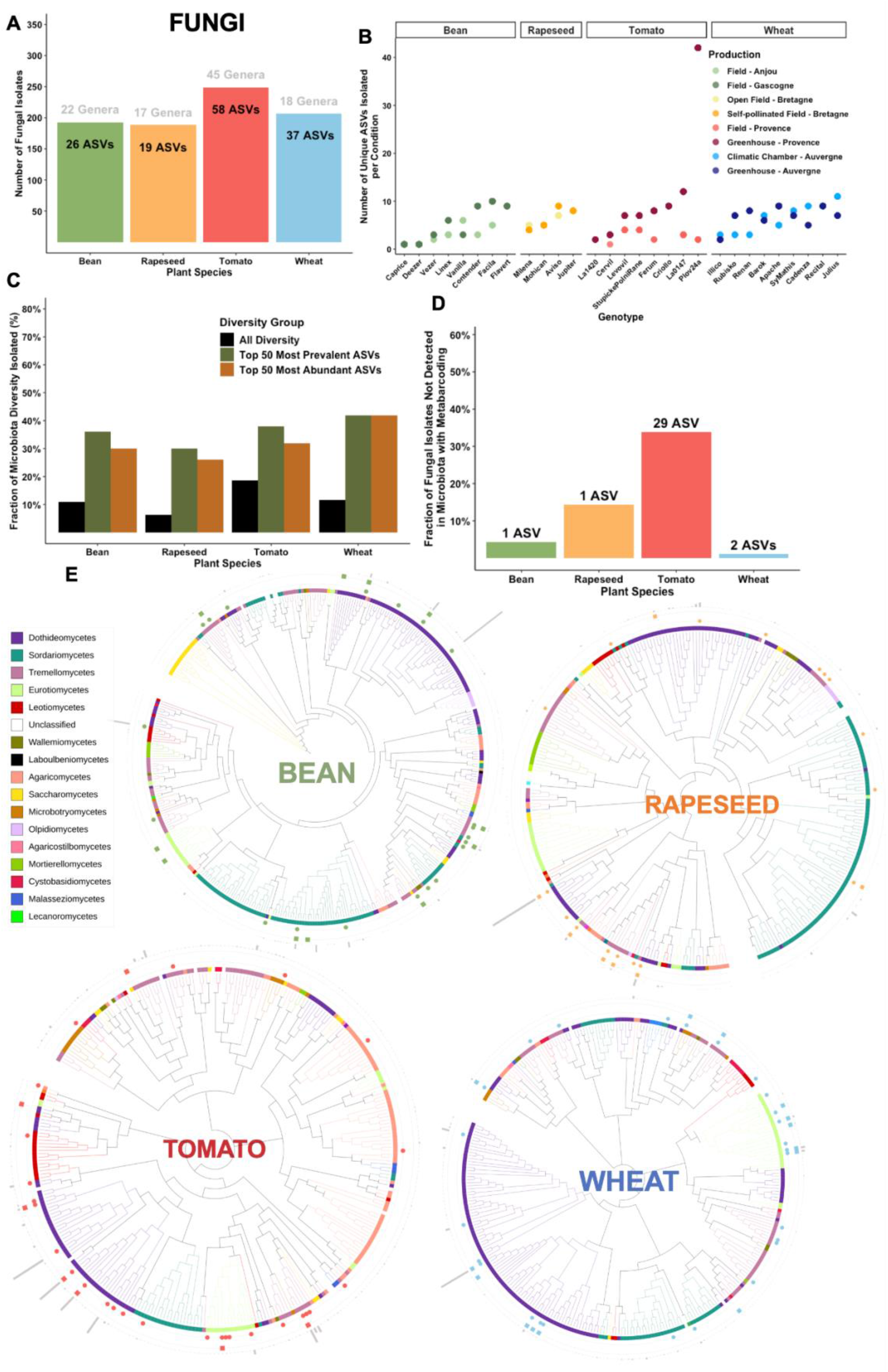
A) Total number of fungal strains isolated by plant species. B) Number of unique strain ASVs isolated for each condition (i.e a genotype in a given production mode) based on both seed and seedling isolates. C. Fraction of seed fungal diversity successfully isolated, based on the comparison of strains ASVs and all ASVs detected in metabarcoding. D) Fraction of strains ASVs only found in the strain collection and not in the metabarcoding analysis. E) Phylogenetic trees of all ITS1 region ASVs detected in seed microbiota based on metabarcoding analysis for each plant species separately. ASVs for which a fungal isolate is available in the collection are indicated by a symbol. The colored inner circle represents the ASV taxonomic affiliation at the phylum level and vertical grey bars on the outer circle represent the average relative abundance of each ASV.

Next, we assessed the phylogenetic representativeness of the strains isolated compared to seed microbiota diversity characterized using metabarcoding (Figure 8E) by representing all the ITS1 ASVs detected on a phylogenetic tree for each plant species. The majority of isolates recovered belonged to *Ascomycota* which is dominant for all plant species, followed by *Basidiomycota* and *Mucoromycetes*. The visualization of the trees confirms the isolation of ASVs with high relative abundance, often from both seeds and seedlings, indicating their ability to transmit to the next generation.

The number of isolates by fungal genus obtained from the four plant species is presented in Figure 9. The culturing effort from seeds was always more important than from seedlings in the four plant species (Figure S3C). Even if the isolation effort by habitat was unbalanced, we can note that isolates from Dothideomycetes and Eurotiomycetes were frequently recovered from seedlings, compared to Tremellomycetes which were mainly isolated from seeds (Figure S3D).

**Figure 9:**
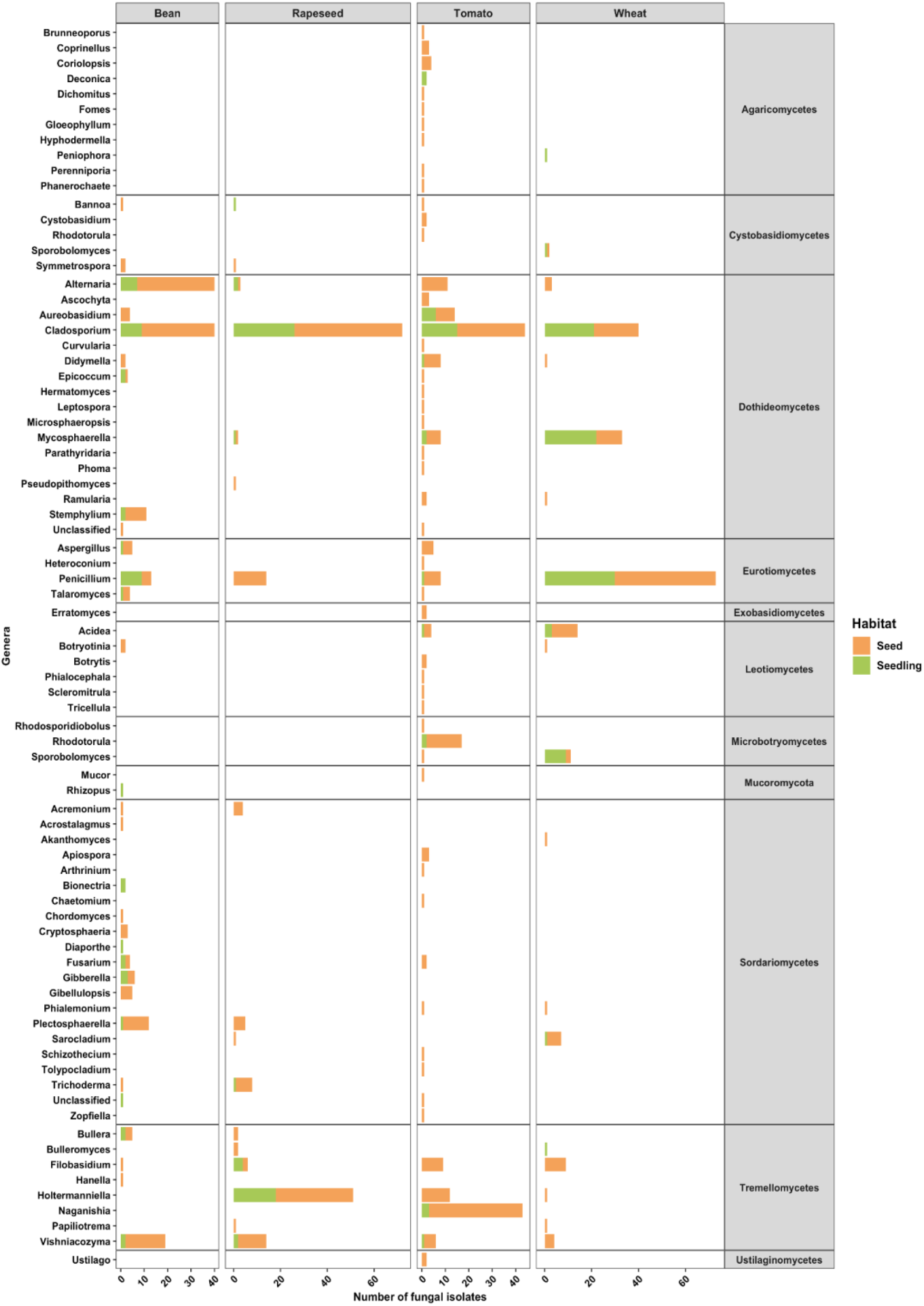
List of all fungal genera isolated from the four plant species. For each genus the number of isolates colored by their source of isolation (seed or seedling germinated *in vitro*) is indicated. The genera are organized by fungal classes.

Isolates affiliated to *Cladosporium, Penicillium, Alternaria, Vishniacozyma* and *Filobasidium* were recovered from all 4 plant species. Out of the 45 shared ASVs between the four species previously identified (Figure 5G), 24 of them were successfully isolated including the 9 most prevalent taxa, such as *Cladosporium perangustum* 1, *Alternaria spp*, or *Epicoccum nigrum*. For the other genera, their isolation varied more between plant species. For instance, *Mycosphaerella, Holtermaniella, Acidea and Aureobasidium* were frequently isolated from two or three species out of the four. A high number of isolates from the *Agaricomycetes* class or *Naganishia* and *Rhodotorula* genera were exclusively isolated from tomato. *Stemphylium* isolates were obtained from bean and *Sporobolomyces* mainly from wheat. These observations are in line with the metabarcoding results showing both shared taxa and distinct compositions between the four plants, with a higher fungal diversity in tomato seeds (Figure 3A, 5).

## DISCUSSION

### Seed Microbiota Variability

In this study, we extend knowledge on the characterization of seed microbiota diversity of four major crops and build a large strain collection of seed-borne bacteria and fungi isolates. Seed microbiota was highly variable between samples and conditions, in terms of bacterial colonization (10 CFUs to 100 million CFUs per g of seed) but also in microbial richness (4 to 351 bacterial ASVs & 16 to 138 fungal ASVs). These results align with previous studies showing high variability in seed microbiota assembly even when produced at the same location over time (Rochefort *et al*. 2019, Morales Moreira, Helgason, and Germida 2021) or from one seed to another (Bintarti *et al*. 2021, Chesneau *et al*. 2022, Kim, Kim, and Lee 2023). An interesting observation was that seeds produced in field environments had higher bacterial colonization than in confined environments (greenhouse or climatic chambers, consistent with Chen *et al*. 2024) but the opposite pattern was observed on microbial richness. Tomato seeds had generally higher bacterial and fungal diversity in the greenhouse production than in the field and similarly wheat seeds were more diverse in the most confined environment of the climatic chamber than in the greenhouse production. This observation confirms that seed-borne microorganisms are in majority acquired from the external environment with a minor role of internal transmission in structuring the overall community (Chesneau et al. 2022, Chen et al. 2024). The higher richness observed under low microbial dispersal in confined environments is surprising but it has been reported in experimental studies a dispersal rate-dependent effect on community diversity, with increasing rates of dispersal resulting in weakened local selection and a hump-shaped pattern of local species diversity (Custer, Bresciani, and Dini-Andreote 2022).

### Divergent Ecological Drivers: Host-Specific Bacteria vs. Stable Fungal Core Members in Seed Microbiota

Another interesting finding was that the bacterial and fungal communities appear to respond to different ecological drivers in seed microbiota. Bacterial communities were more host-specific and variable, while fungal communities were more stable and dominated by a core set of taxa shared across different plants. Fungi presented a more substantial core community across all plant species (45 shared ASVs contributing 68.79% of abundance), whereas bacteria had fewer shared ASVs across all plants (21 ASVs contributing 25.5% of abundance). These results suggest that bacterial community assemblies are more driven by stochastic processes (e.g dispersal, drift), while fungal communities are under more deterministic processes (e.g host and environmental selection). The core shared taxa were *Pantoea agglomerans, Variovorax sp, Rhizobium sp, Cutibacterium sp* or *Sphingomonas sp* for bacteria, *Cladosporium perangustum, Alternaria* (*chromatica &* sp), *Epicoccum nigrum* for filamentous fungi and *Sporobolomyces roseus* and *Vishniacozyma victoriae as* Basidiomycota yeasts. The identification of these shared taxa is in line with a meta-analysis which also identified these taxa as members of the core seed microbiome across 50 plant species (Simonin *et al*. 2022) confirming the existence of highly adapted microorganisms to the seed environment that are widespread in diverse terrestrial ecosystems.

### Genotype and Production effects: Each Seed Sample Produced has a Distinct Microbiota

In this study, we also assessed the influence of host genotype and production mode for each crop independently. While the differences in production modes were not comparable for the different crops (field vs greenhouse or 2 fields in different regions), it was interesting to see that globally seed microbiota was significantly influenced by genotype, production and their interactions in all cases. This means that each seed sample produced under given environmental conditions has a distinct microbiota which can have important implications on seed and plant health. These significant effects of host genotype and production are coherent with the existing literature on diverse crops (Klaedtke *et al*. 2016, Rochefort *et al*. 2019, Malacrinò *et al*. 2023, Tabassum *et al*. 2024). These results suggest that genotype-specific selection occurring during flowering and seed development interacts with external environmental conditions to influence microbiota assembly. Currently, very few studies are available on the assembly mechanisms and transmission pathways of seed-borne microorganisms (i.e internal, floral, external) but these processes deserve more investigation to better understand the variability in seed microbiota structuration (Chesneau *et al*. 2022, Chen *et al*. 2024, Bergmann *et al*. 2025).

### Crop-Specific Patterns in Seed Microbiota

The four crops had clearly distinct seed microbiota colonization and structure for both bacteria and fungal communities. In particular, bean seeds were characterized by a lower bacterial colonization (3.85 log CFU/g seed) despite their large sizes, a high microbial richness and variability in composition. These results are consistent with bean single seed microbiota analysis which harbor very variable composition from one seed to another that can lead to high diversity levels when assessed on a pool of 30 seeds (Chesneau *et al*. 2022).

Rapeseed seeds were strongly influenced by production mode with a higher bacterial and fungal ASV richness observed in the self-pollinated field for the four genotypes but a lower bacterial colonization. The self-pollinated condition relied on fly pollination in insect proof cages to avoid cross-pollination between genotypes, which created more confined conditions compared to open field conditions. These results show the importance of insect pollination on seed microbiota assembly that can introduce diverse taxa through the floral pathway (Prado *et al*. 2020). For instance, the bacterial taxa *Sphingomonas* or *Rhizobium* and fungal taxa *Gibberella* or *Slerotinia* were only present under fly pollination which can be plant pathogens. Tomato seeds were characterized by a higher bacterial colonization and very distinct composition compared to the three other species, likely due to differences in fruit types (fleshy berry fruit for tomato vs dry fruit for the others). Specific signatures of tomato seeds were the dominance of *Acinetobacter* bacteria across all conditions and the presence of *Naganishia* yeasts in field samples. *Acinetobacter* has already been reported as the most abundant member of the tomato phyllosphere and to be highly abundant in fruit placenta, which could explain its dominance in seeds (Dong *et al*. 2019). Naganishia yeasts have been previously isolated from flowers (Zhou *et al*. 2020).

For wheat seeds, the genotype had a large impact on both bacterial and fungal communities in terms of bacterial population sizes, composition and richness. The most dominant taxa were *Pantoea agglomerans, Cladosporium perangustum* and *Alternaria sp* that are known core seed and stem microorganisms with evidences of vertical transmission in wheat (Sharma *et al*. 2023, Sun, Sharon, and Sharon 2023, Becker and Cubeta 2024, Sanz-Puente *et al*. 2025). Altogether the seed material produced in this study yielded a resource of diverse microorganisms, some being shared across crops and others specific to a given genotype or production for our culturomics efforts.

### Microbial Strain Isolation & representativeness

Our culturomics approach yielded a collection of 2,510 bacterial and 837 fungal isolates from seeds of the four plant species. This extensive isolation effort provides valuable insights into the culturable fraction of seed microbiota and its representativeness across different plant hosts. The bacterial diversity of unique isolates recovered was consistent with the sampling efforts, as reflected by the number of colonies picked per sample and the observed bacterial richness. Our approach, based on visual selection of colonies, proved efficient in avoiding the isolation of multiple clones of the same microorganism, thus maximizing the capture of distinct taxa. A significant outcome of our study was the successful isolation of shared taxa across multiple plant species, which align with core members identified by metabarcoding.

The frequency of isolation for different bacterial genera varied among plant species, confirming potential host preferences or adaptations. While some genera like *Pantoea, Curtobacterium, Frigoribacterium, Paenibacillus*, and *Rathayibacter* were commonly isolated from bean, rapeseed, and wheat, they were generally absent from tomato samples. Conversely, *Acinetobacter* and *Massilia* were more frequently isolated from tomato seeds, in line with metabarcoding results. Our findings also align with the meta-analysis of culturable plant bacterial endophytes by Riva et al., which identified a culturable core including *Bacillus, Pantoea, Pseudomonas, Microbacterium, Paenibacillus*, and *Enterobacter*. Also consistent with this meta-analysis, our study reveals that the majority of the isolated taxa were rare (just few isolates), highlighting the importance of extensive sampling to capture the diversity of culturable seed microbiota. It is noteworthy that this level of representativeness was achieved using only one medium (TSA 10% strength), previously described as a generalist medium for plant bacteria (Chesneau *et al*. 2022, Zhang *et al*. 2021). This suggests that our approach was efficient in capturing a broad spectrum of culturable bacteria from seed microbiota. However, we acknowledge that even higher representativeness could be achieved through the use of plant-based media (Sarhan *et al*. 2019), or by employing advanced culturomics technologies such as automated microbiome imaging and isolation (Huang *et al*. 2023).

Our culturomics approach also enabled the recovery of a substantial fungal collection, spanning a wide diversity of taxa across the four plant species. As observed for bacteria, the diversity of fungal isolates was consistent with metabarcoding-based richness patterns, with tomato seeds harboring the highest diversity and rapeseed the lowest. Despite recovering only 6 to 18% of the total ASV richness, our collection captured a large fraction (74 to 82%) of the relative abundance of the seed mycobiota, indicating that dominant and ecologically relevant taxa were efficiently isolated. These numbers are consistent with a previous study on the isolation of fungal endophytes which recovered 15% of the diversity detected by metabarcoding (Durán, San Emeterio, and Canals 2021). Approximately one third of the most prevalent and abundant fungal taxa were successfully isolated across all plant species, including core taxa such as Cladosporium, Alternaria, and Epicoccum, supporting the relevance of this collection for downstream functional analyses. Phylogenetic analysis further confirmed that the isolates broadly recapitulate the taxonomic structure of the seed mycobiota, with a strong representation of Ascomycota and, to a lesser extent, Basidiomycota and Mucoromycetes. In addition, the recovery of isolates from both seeds and seedlings for several abundant ASVs suggests effective vertical transmission and reinforces their ecological importance. Finally, the relatively low proportion of fungal isolates undetected by metabarcoding, except in tomato, supports the overall congruence between approaches.

Future methodological enhancements for culturomics could further bridge the gap between culturable and total microbial diversity in seed microbiota, providing a more comprehensive understanding of these highly variable microbial communities. Future studies should focus on the identification of additional rare or fastidious microorganisms contributing to seed ecology and health.

### Culturomics vs. Metabarcoding

Our study highlights the complementary nature of culturomics and metabarcoding in characterizing seed microbiota. While metabarcoding provides a broad overview of microbial community composition, culturomics enables the detection of rare bacterial taxa that may be underrepresented or entirely missed in sequencing-based approaches due to low PCR amplification efficiency (Laval *et al*. 2021, Moinard *et al*. 2023). This finding is consistent with previous plant culturomics studies by Chesneau *et al*. (2022) and Ren *et al*., (2023.), which demonstrated that culturomics can recover a large fraction of microbial diversity undetected by metabarcoding, offering a more comprehensive view of microbial diversity. By integrating these two approaches, researchers can achieve a more holistic understanding of seed microbiota, capturing both the dominant and rare microbial members and their potential ecological roles.

## Conclusion

This study demonstrated that each seed sample produced harbors a specific assemblage of bacteria and fungi, reflecting the influence of both host-driven and environmental factors on seed microbiota composition. The acquisition of microbes from the environment means that seed production under controlled conditions, such as greenhouses or climatic chambers, may not fully represent native microbial communities, often resulting in lower microbial biomass but increased diversity. We also observed that fungal communities tend to be more stable than bacterial assemblages with a significant fraction shared across genotypes and production systems. The fact that bacteria and fungi seem to be structured by distinct ecological drivers is an important finding that deserves more attention for the successful development of seed microbiota engineering in agriculture.

Culturomics has proven to be an efficient approach for studying seed microbiota, allowing the isolation of a large fraction of highly abundant and prevalent taxa while also enabling the identification of rare taxa that may be overlooked by metabarcoding methods. The isolation of core and host-specific taxa offers new avenues for investigating the functional roles of these microorganisms in seed biology and plant health. Together, these findings highlight the complexity and variability of seed microbiota, emphasizing the need for integrative approaches to better understand their roles in plant health and productivity.

## Acknowledgements

This work was funded by the 3rd Programme for Future Investments (France2030) and operated by the SUCSEED project (ANR-20-PCPA-0009) funded by the ‘Growing and Protecting crops Differently’ French Priority Research Program (PPR-CPA), part of the national investment plan operated by the French National Research Agency (ANR). This research was also funded through the OSMOSE project in the framework of the regional programme ‘Objectif Végétal, Research, Education and Innovation in Pays de la Loire’, supported by the French Region Pays de la Loire, Angers Loire Métropole and the European Regional Development Fund. We thank Anne-Sophie Poisson for her work managing the SUCSEED project. We thank Elise Alix and Bernard Moulin (IGEPP) and the Experimental Unit UE La Motte for the rapeseed seed production in the field and the CRB BrACYSol for providing the rapeseed genetic resources. We thank Muriel Bahut (ANAN platform, SFR Quasav) for amplicon sequencing. We are thankful to Mélanie Andrin for taking care of the plants, to the vegetable resources centre (CRBLeg) of GAFL for keeping the seeds available, and to the staff of UE A2M ‘Arboriculture et Maraichage Méditerranéens’. We thank the FNAMS (Fédération Nationale des Agriculteurs Multiplicateurs de Semences) for producing the bean seeds at two sites (Brain sur l’Authion and Condom).

## SUPPLEMENTARY MATERIAL

**Figure S1:**
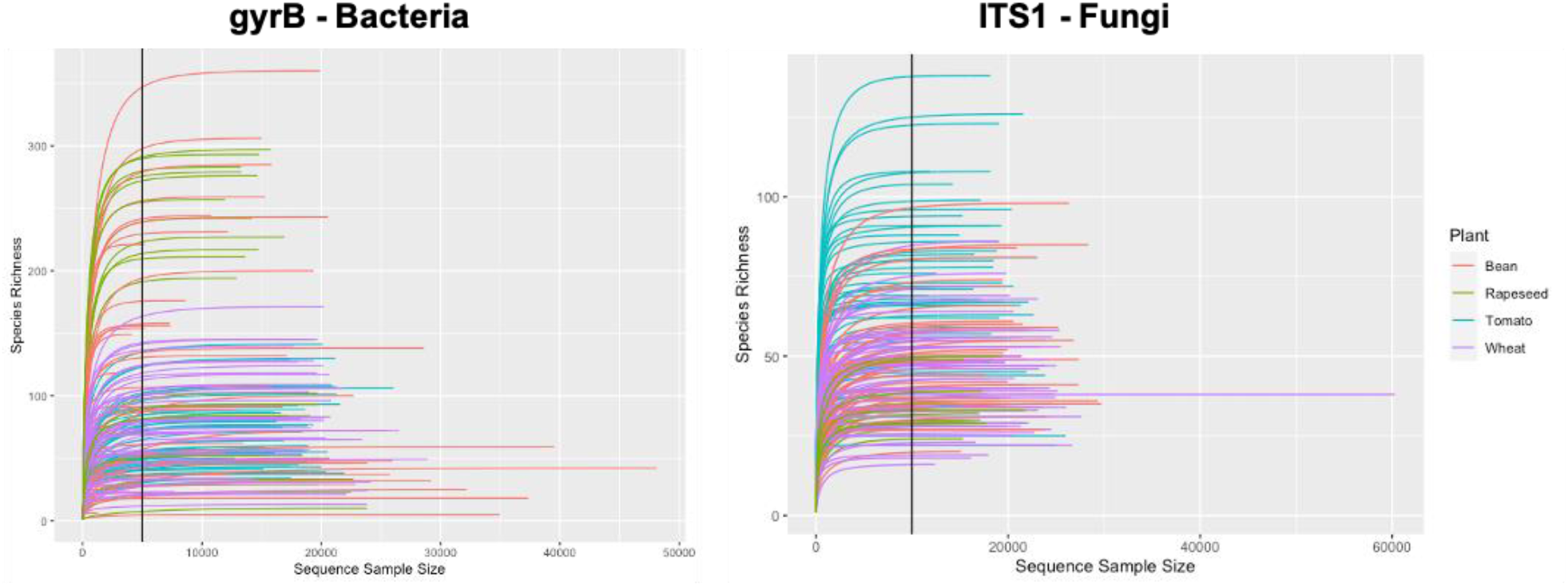
Rarefaction curves for the gyrB (Bacteria) and ITS1 (Fungi) marker genes datasets. The vertical line indicates the rarefaction level chosen for the alpha diversity analysis. Samples with lower read counts were excluded from the analysis.

**Figure S2:**
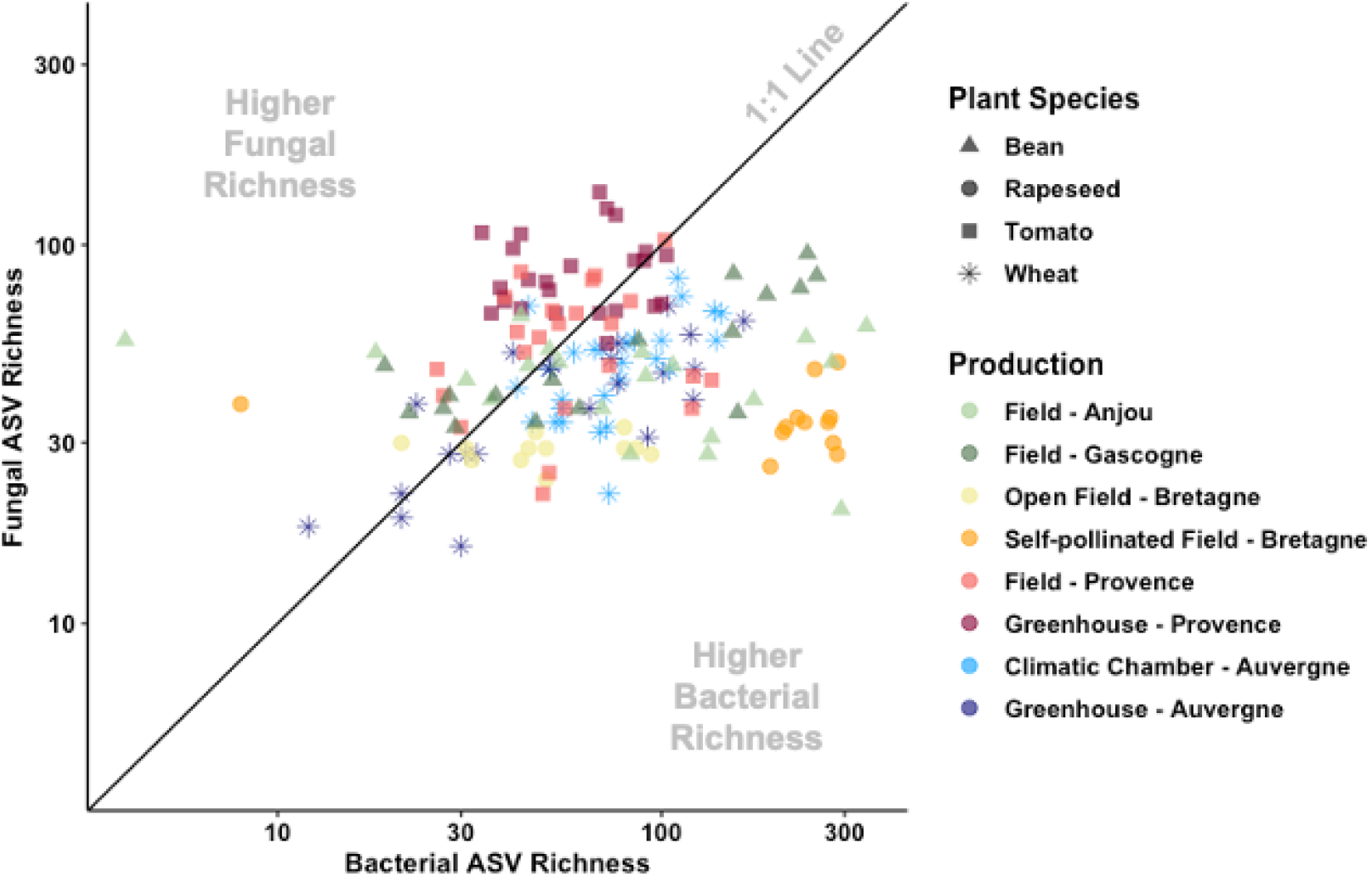
Relationship between bacterial and fungal observed richness ratio (expressed per seed sample). The 1 : 1 line facilitates the visualization of samples presenting an enrichment of one of the taxonomic groups over another. Note that both axes are on a log-scale.

**Figure S3:**
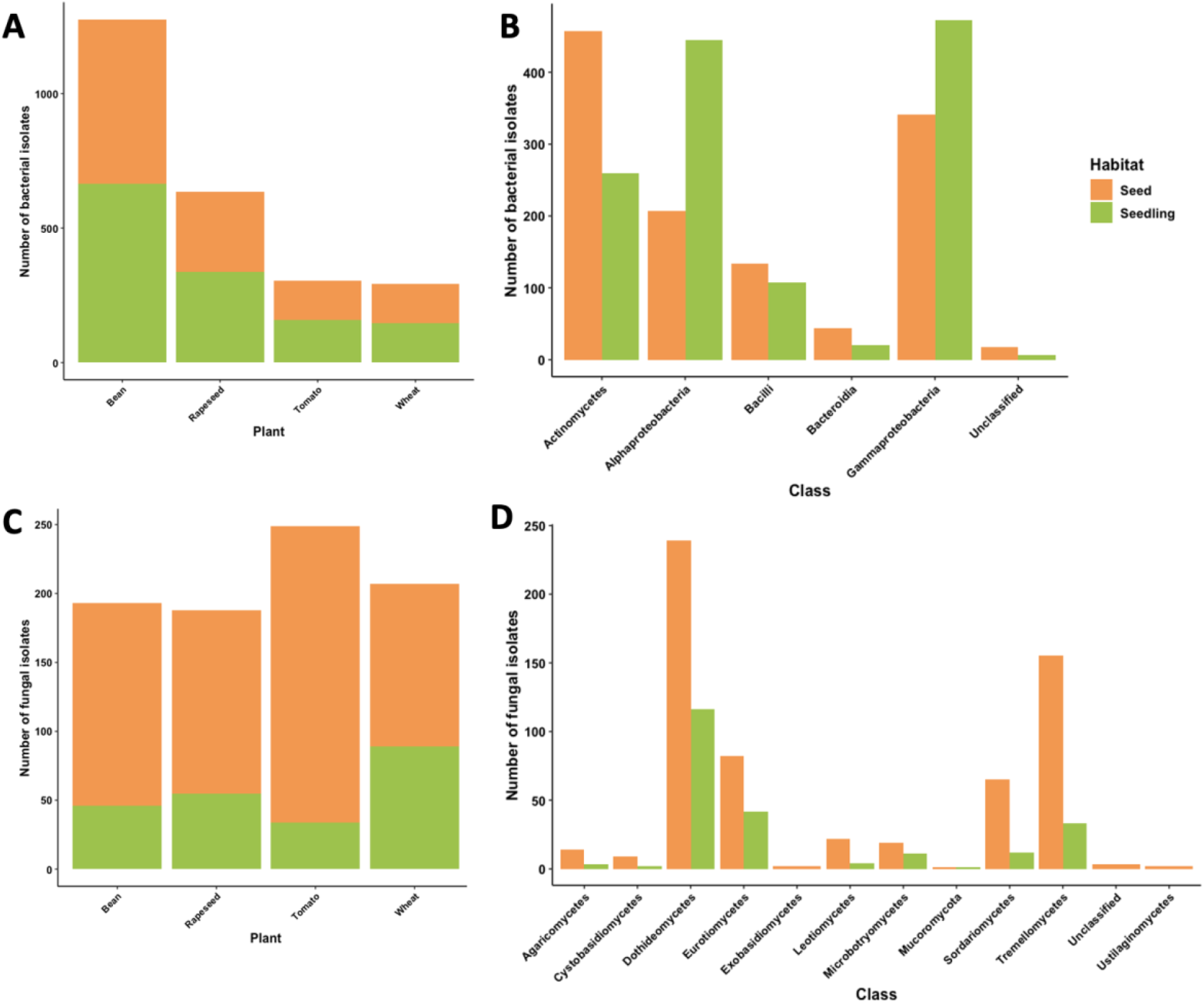
Number of bacterial isolates recovered from seeds or seedlings germinated in vitro (sterile conditions), by A) plant species and B) by bacterial classes. Number of fungal isolates by C) plant species and D) by fungal classes.

